# Essential Role of MHC II in the Antitubercular Efficacy of Pyrazinamide

**DOI:** 10.1101/2025.08.21.671522

**Authors:** Elise A. Lamont, Shannon L. Kordus, Michael D. Howe, Ziyi Jia, Nathan Schacht, Muzafar Rather, Gebremichal Gebretsadik, Anthony D. Baughn

## Abstract

Antibacterial drug mechanisms have traditionally been examined through a drug-pathogen lens, often overlooking the host’s role in shaping drug activity. However, growing evidence suggests that the host environment is crucial for antibacterial efficacy. Pyrazinamide (PZA), a key component of modern tuberculosis therapy, exemplifies this complexity—exhibiting potent *in vivo* activity despite its inability to reduce *Mycobacterium tuberculosis* viability in standard *in vitro* culture. Here, using macrophage and murine infection models, we identify a critical role for host cell-mediated immunity in PZA’s antitubercular action. Through the use of MHC II knockout mice, we demonstrate that CD4 T cell help is essential for PZA efficacy. Notably, while IFN-γ is required for PZA-mediated clearance of *M. tuberculosis* at extrapulmonary sites, bacterial reduction in the lungs occurs independently of IFN-γ signaling. Additionally, we show that PZA leverages cell-mediated immunity in part through activation of the oxidative burst. Our findings underscore the need to incorporate host factors into antibacterial drug evaluation and highlight potential avenues for host-directed therapies and adjunctive antibiotics in first- and second-line tuberculosis treatment.

## Introduction

“…when pyrazinamide is used alone, we are never entirely certain what will happen,” Walsh McDermott, a pioneer of multidrug chemotherapy for tuberculosis (TB), exclaimed in his 1956 paper promoting a pyrazinamide (PZA) and isoniazid (INH) combinatorial regimen (1). This sentiment was shared by McDermott’s contemporaries in the TB field and often led to finger pointing at the host for the cause of uncertainty in drug activity, particularly when treatment failure could not be attributed to drug resistance (1). The recognition of the host environment being a significant contributor to drug activity resulted in defining drug therapy as a three-way relationship involving the drug, parasite, and host (1, 2). In fact, the initial discovery of PZA as a sterilizing drug to treat TB was made possible by *in vivo* studies in *Mycobacterium tuberculosis* (*Mtb*) infected mice (3, 4). Under standard *in vitro* broth and agar conditions, PZA displayed no discernable growth effects against *Mtb* cultures (5-7). Thus, the host environment was considered a fertile basis for uncovering drug mechanisms and activities, especially for drugs that may have significantly disparate responses under *in vitro* and *in vivo* conditions (2).

Throughout the evolution of clinical drug use, the mechanism of drug action has largely been untethered from the host response partially due to the complexity and reproducibility of pharmacodynamic studies in animal models and the patient care setting (8, 9). Within the last 18 years, there has been a modest pivot back to the host as a handful of antibiotics were found to influence the immune response such as antibiotic induced neutrophil extracellular traps and secondary responses such as reactive oxygen species (ROS) production as a byproduct of altering bacterial metabolism (10-12). However, these findings illustrate antibiotics acting on the host rather than the host as a potential influencer of antibiotic action and success. Ultimately, drug activity with regards to discovery and optimization has taken a reductionist approach and centered on defining the drug-parasite relationship in terms of the drug’s minimum inhibitory concentration (MIC) using standardized bacteriologic media (13). Bacteriologic media can be supplemented with purified antimicrobial peptides and small chemicals to mimic aspects of the host response for drug testing. Indeed, this has been the case for PZA testing and has identified several stressors that promote and enhance its activity, such as low pH, nutrient limitation, and hypoxia (5, 14, 15). However, the full dynamics of the host environment and its impact on drug activity and localization cannot fully be appreciated using *in vitro* cultures alone. To date, PZA remains the only drug in the current TB pharmacopeia in clinical use that has an unknown mode of action (16). Several models have been proposed to explain PZA’s mechanism of which only one (the PZA protonophore model) describes the host as a contributor in PZA activity (16-18). Although PZA as a protonophore has been challenged, acidic pH remains an integral component of host-mediated potentiation of PZA and an essential piece in understanding how PZA functions *in vivo* (19, 20).

In recent years a growing body of evidence encompassing both animal models and human granuloma specimens have highlighted the critical importance of the host environment on PZA penetration, availability, and activity (21-28). Several studies have demonstrated that *Mtb* susceptibility to PZA is influenced by the location of the tubercule bacilli within the granuloma and individual cells (23, 25, 29). In cell models of TB infection, PZA was shown to significantly accumulate in acidified phagosomes and thereby produce maximum efficacy (20, 29). Furthermore, several studies have reported diminished or abrogated PZA activity in mouse strains with compromised cell-mediated immunity (23, 24, 26). Additionally, this research has shown that PZA fails to sterilize extracellular *Mtb* bacilli residing in the near neutral pH caseum of the C3HeB/FeJ mouse strain (25). Caseum pH appears to be an important aspect of this particular host environment as purified acidic caseum from a rabbit model showed modest cidal activity by PZA (30, 31). Although PZA-mediated killing of extracellular *Mtb* can be achieved in caseum at an acidic pH, PZA is likely most potent against *Mtb* localized to intracellular environments, like the phagosome.

Within the intracellular environment, several host defense mechanisms are initiated against *Mtb*. Some of these host responses have been associated with PZA potentiation and it is possible that other related stressors are yet to be identified. Elucidating which host responses that synergize or bioactivate PZA will guide adjunct host directed therapies that shorten the current TB regimen and overcome TB drug failure in some patient populations.

We have previously shown that *Mtb* bacilli exposed to sublethal concentrations of PZA are sensitized to reactive oxygen species (ROS), resulting in enhanced bactericidal activity (32). Host derived ROS via NADPH oxidase upregulation by gamma-interferon (IFN-γ) is a critical defense strategy to restrict *Mtb* growth, which culminates in the oxidative burst for pathogen elimination (33-35). We have also shown that IFN-γ activation of macrophage cell lines increased PZA killing of *Mtb* (32). These data suggest that ROS is another exciting host related component of PZA mechanism and may explain the basis for PZA’s conditional susceptibility *in vivo*. In this study, we further investigate the role of ROS, the oxidative burst, and the cell-mediated immune response in PZA potentiation. Using complementary cell and animal models of TB infection, we show that the oxidative burst and MHC II expression are essential for the potent bactericidal activity of PZA *in vivo*. This previously unknown mechanism will allow for the design and inclusion of host-directed therapies that will initiate and amplify the oxidative burst to promote PZA function.

## Results

We selected bone marrow derived macrophages (BMDMs) from well-established genetic knockout mouse strains (gamma-interferon receptor 1 (*ifngR1*; deficient in gamma-interferon signaling) KO and *phox47* KO (deficient in producing p47phox) and wild-type counterparts (C57BL/6J and C57BL/6NJ) in *Mtb* infection assays to further explore the role of the oxidative burst in PZA activity. Previous studies from our group have shown that ROS sensitizes *Mtb* bacilli to PZA and thus enhances drug activity (32). To promote the oxidative burst, we activated BMDMs with IFN-γ prior to PZA treatment. IFN-γ upregulates NADPH oxidase, a multi-subunit enzyme complex that catalyzes molecular oxygen to generate superoxide anions, which are later converted to multiple ROS within the phagosome (35, 36). Bacterial burden was assessed throughout the 5-day post-infection (p.i.) period. Similar to our previous findings in *Mtb* infected RAW 264.7 and THP-1 macrophage-like cell lines, IFN-γ activated BMDMs enhanced PZA-mediated killing of *Mtb* in both C57BL/6J and C57BL/6NJ WT BMDMs (32). The addition of IFN-γ to PZA (200 µg/mL) resulted in a 2.65- and 1.51-log_10_ reduction in bacterial burden in C57BL/6J and C57BL/6NJ BMDMs, respectively, greater than PZA (200 µg/mL) alone at 5 days p.i. (Fig. 1A and C, Tables S1 and S2). Increased bacterial killing was also observed at days 4 and 5 p.i. in the IFN-γ and PZA (400 µg/mL) treatment compared to PZA (400 µg/mL) alone in BMDMs from both WT strains of mice (Fig. 1A and C, Tables S1 and S2). This elevated bactericidal activity is due to the synergy between IFN-γ and PZA as this combinatorial treatment resulted in a 0.5-1.4 greater log_10_ reduction than IFN-γ alone at day 5 p.i. for both WT strains (Fig. 1A and C, Tables S1 and S2).

**Figure 1.**
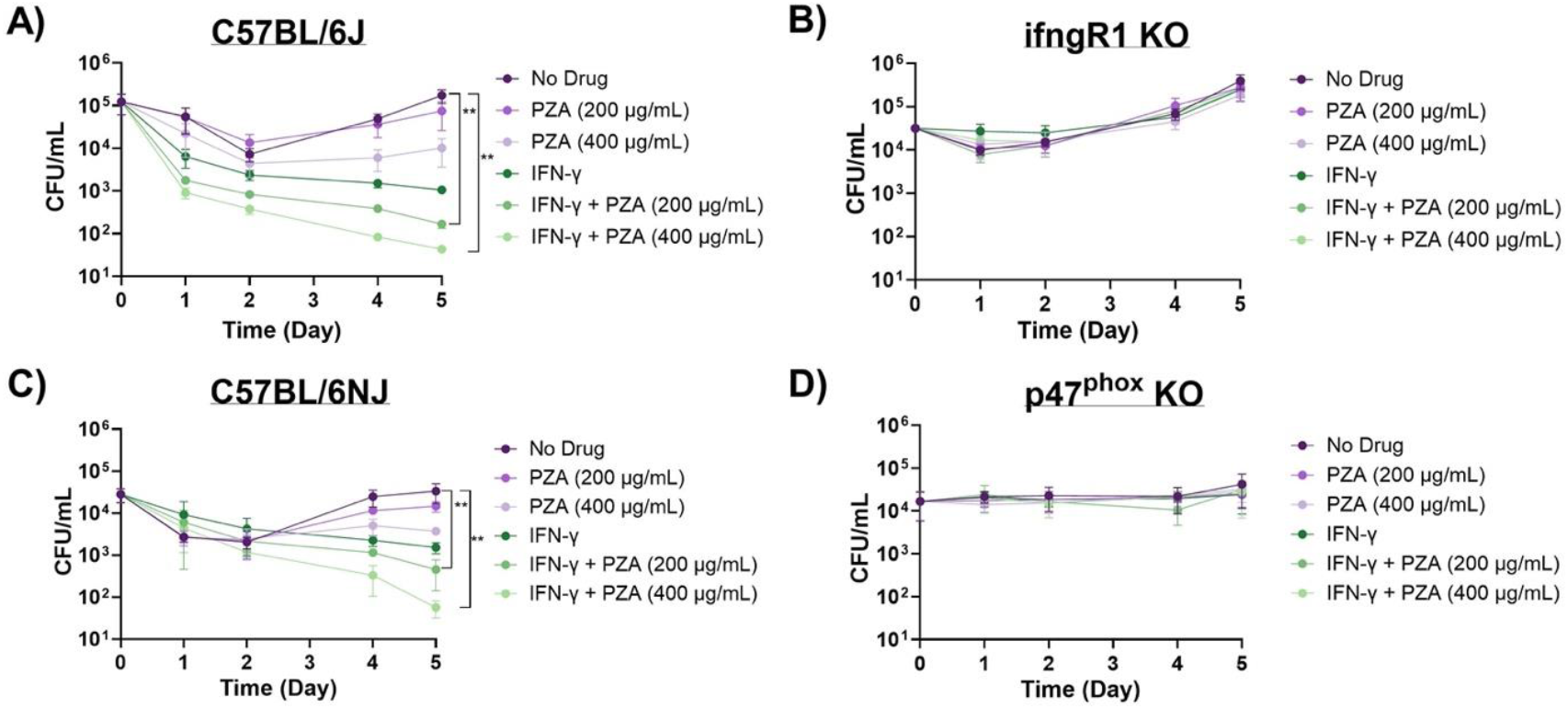
PZA activity is dependent on the oxidative burst. PZA bactericidal activity in *Mtb* strain H37Rv infected (A) C57BL/6J, (B) ifngR1 KO, (C) C57BL/6JN, and (D) p47^phox^ KO BMDMs. Infected resting and IFN-γ activated BMDMs were untreated or treated with PZA (200 µg/mL or 400 µg/mL). Cultures were plated for bacterial enumeration at indicated timepoints. Statistical significance was calculated based on multiple comparisons with 2-way ANOVA with Bonferoni correction. \*\**P<0*.*005*.

IFN-γ and PZA synergy was abrogated in *ifngR1* KO BMDMs and was indistinguishable from the no drug control (Fig. 1B). IFN-γ was unable to bind to IFNGR1; therefore, preventing further signaling including increasing the oxidative burst capacity. While IFN-γ is critical to the oxidative burst, it also regulates pro- and anti-inflammatory cytokines including TNF-α, IL-1, IL-6, IL-12, and IL-10 (37-39). It is possible that IFN-γ enhancement of PZA is due to the modification of these cytokines that are unrelated to the oxidative burst. For example, unique among the first-line TB drugs, PZA was shown to be active in highly inflamed lesions during early treatment and may explain why this drug is at its most potent in the first two months of therapy (27, 40). More recently, short-term blocking of IL-10 using anti-IL-10R1 antibody in *Mtb* infected mice aided PZA-mediated bacterial clearance in 70% of treated mice (below the threshold of detection) compared to mice given PZA alone within 30 days posttreatment (41). Given the above, *phox47* KO BMDMs were subsequently used to directly assess the oxidative burst in PZA potentiation. p47^Phox^ is a critical regulatory subunit in the NADPH oxidase complex, which aids in ROS production (42). A previous study demonstrated that *p47*^*phox-/-*^ mice failed to control *Mtb* replication during early infection and that this loss of control was most likely attributed to diminished ROS production (43). Our results show that the addition of IFN-γ is unable to enhance the oxidative burst in BMDMs from *phox47* KO mice and consequently fails to alter PZA activity against *Mtb* (Fig. 1D). Together these cell infection data suggest that the oxidative burst is an important host component for PZA-mediated killing of *Mtb*.

Previous studies have demonstrated a loss of PZA activity in athymic nude mice but have not further delineated which host component is responsible for this stunning observation (24). In fact, multiple *in vivio* studies have observed refractory responses from PZA treatment in mice deficient in cell-mediated immunity (21, 23-26). Based on these reported findings and our own results highlighting the contribution of IFN-γ and the oxidative burst in PZA activity, we asked what the role of IFN-γ in PZA efficacy is in vivo. *Mtb* infected C57BL/6J (WT) and *ifngR1* KO mice were treated with controls (vehicle control (sterile water) or INH) or PZA 3-weeks p.i. commensurate with the initiation of cell-mediated immunity. Treatments were administered via daily oral gavage for 2-weeks and lung, spleen, and liver bacterial burdens were assessed. As expected, PZA treatment in WT mice caused a reduction in bacterial load compared to vehicle control for all organs (Fig. 2). PZA treatment reduced *Mtb* bacterial load by 2-, 1.5-, and 1.5-log_10_ in lungs, spleens, and livers, respectively, compared to the vehicle control (Fig. 2). However, PZA treatment in *ifngR1* KO mice displayed disparate responses in the lungs versus extrapulmonary sites (Fig. 2). In the lungs of *ifngR1* KO mice, PZA-mediated killing of *Mtb* remained intact albeit to a lesser extent (∼1.2-log_10_ reduction in bacterial burden compared to 2-log_10_ reduction in WT counterparts; Fig. 2). However, in both *ifngR1* KO spleens and livers PZA treatment was indiscernible from the vehicle control (Fig. 2). Drug failure in the *ifngR1* KO mice spleens and livers are specific to PZA as the control drug, INH (structurally similar to PZA), results in 2.5- and 1-log_10_ reductions in *Mtb* bacterial loads in spleens and livers, respectively (Fig. 2). In fact, some *ifngR1* KO mice treated with INH had no discernable bacterial growth in the spleens and livers (below limit of detection; Fig. 2). The discrepancy in PZA-mediated *Mtb* killing may partially be explained by recent findings demonstrating PD-1 repression of IFN-γ in the lungs to control host-induced immunopathology (44). In contrast, over 80% of bacterial replication is restricted by IFN-γ in the spleen (44). Thus, IFN-γ is likely to be a significant contributor of PZA potentiation at extrapulmonary sites where increased IFN-γ production is not detrimental to the host.

**Figure 2.**
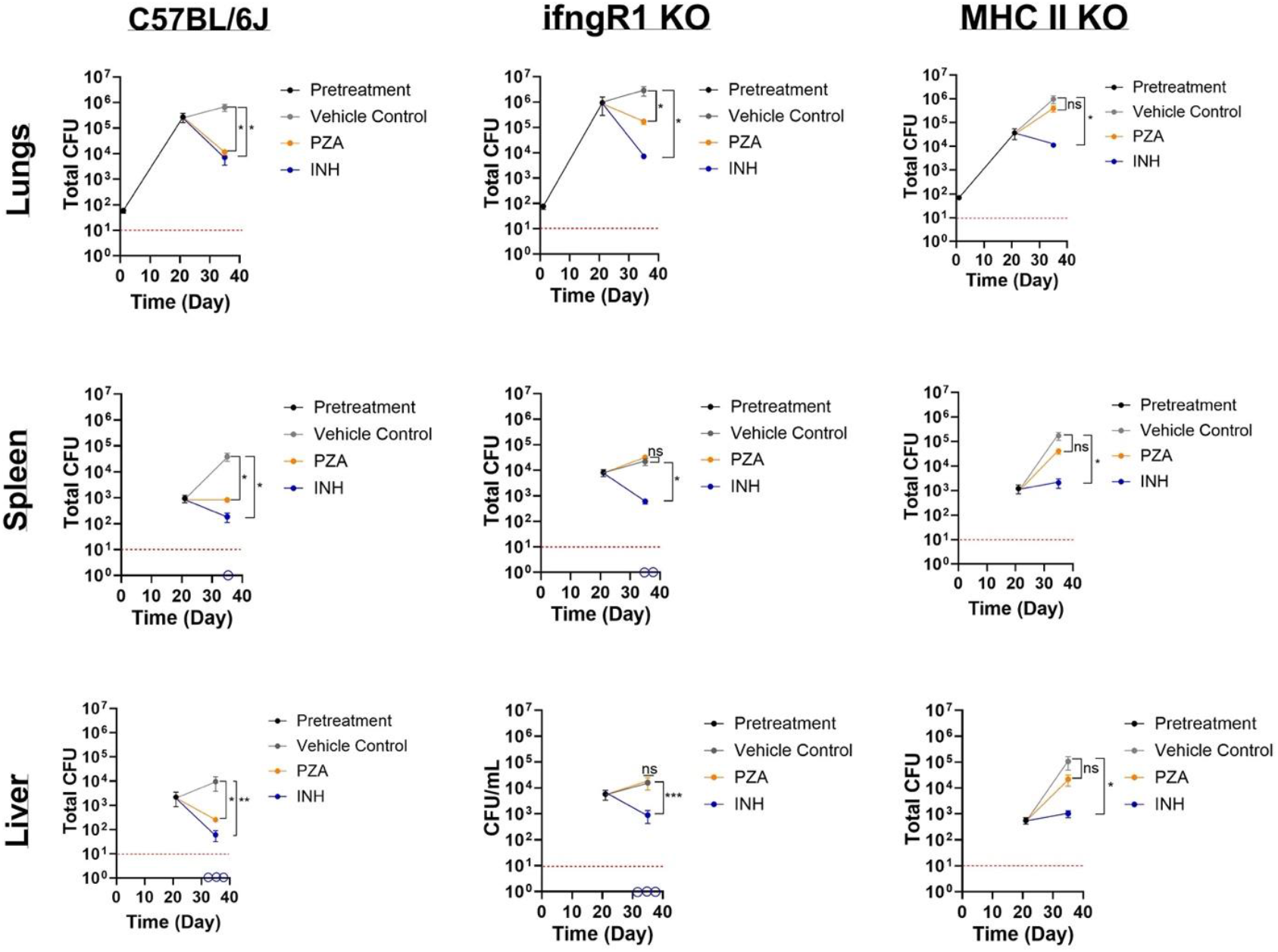
PZA failure in ifngR1 KO and MCH II KO mice. *Mtb* strain H37Rv total bacterial burden in lungs, livers, and spleens of C57BL/6J, ifngR1 KO, and MHC II KO mice. All mice were infected with low-dose aerosol *Mtb* and the infection was allowed to progress for 3-weeks in order for cell-mediated immunity to initiate (pretreatment). Mice were subsequently treated daily for 5 days a week for 2 weeks with either vehicle control (sterile water), PZA (150 mg/kg), and IN (30 mg/kg). Upon euthanasia, organs were removed, homogenized, and plated for bacterial enumeration at indicated timepoints. Empty circles on the x-axis indicates animals for which no bacterial growth was detected. Statistical significance was calculated based on paired two-tailed *t*-tests and *p* values were adjusted for multiple comparisons.*\*P*<0.05, *\*\*\*P*<0.0005, ns not significant.

IFN-γ independent mechanisms exist within the lung and serve as significant contributors in the host response to pulmonary TB. For instance, CD4 T cells expressing CD153 (a tumor-necrosis factor (TNF) super family member) have been correlated with reduced lung *Mtb* load in non-human primates and humans independent of IFN-γ production (45). Furthermore, CD153 was shown to be critical in the control of *Mtb* replication in the lungs of infected mice (44). It should also be noted that CD153 is a ligand of CD30, a co-stimulatory receptor that was revealed to be necessary for CD4 T cell control of *Mtb* expansion in the lung (46). Together, both receptor (CD30) and ligand (CD153) drive CD4 T cell differentiation and multiple effector responses. Given the importance of CD4 T cells in pulmonary *Mtb* control combined with IFN-γ independence in some cell subsets, we proceeded to ask if CD4 T cells played a role in PZA activity in the lung. We selected MHC class II (MHC II) KO mice to pursue this question as this mouse strain is deficient in MHC II and lacks CD4^+^ T cells (47). *Mtb* infection and drug treatment in MHC II KO mice was performed exactly as the previous *in vivo* experiment. Regardless of tissue type, PZA failed to make an appreciable difference in *Mtb* burden in MHC II KO mice (Fig. 2). Unlike the ifngR1 KO mice, PZA failed to reduce the number of *Mtb* bacilli from the recorded pre-treatment value (Fig. 2). While slight differences in *Mtb* load were noted for the spleen and liver (∼0.4 and ∼0.52-log_10_ decreases, respectively), these reductions were not statistically different than the vehicle control (Fig. 2). Treatment failure in MHC II KO mice appears to be restricted to PZA as INH reduced bacterial burden by 2-log_10_ in lungs and extrapulmonary organs (Fig. 2). These data suggest that the cell-mediated component of PZA potentiation is MHC II dependent.

## Discussion

To conclude, this is the first study to specifically demonstrate MHC II expression, a strong inference for CD4 T cells, as a major host factor responsible for the conditional susceptibility of PZA *in vivo*. Both IFN-γ dependent and independent CD4 T cells are likely to play a key function in PZA activity that is specialized to host tissues. Based on our findings, IFNγ^+^ CD4 T cells are anticipated to promote PZA action in extrapulmonary organs, while IFNγ independent CD4 T cells contribute to PZA-mediated *Mtb* clearance within the lung. Future studies should further explore the contribution of specific subsets of CD4 T cells to PZA function. Particularly, CD30 KO and CD153 KO mice should be utilized to assess their contribution to PZA activity in the lungs. Elucidating CD30’s role in PZA-mediated *Mtb* clearance will also be essential to inform drug therapy decisions in TB positive patients that also have relapsed or refractory lymphoma. Relapsed/refractory lymphoma patients may be prescribed anti-CD30 monoclonal antibody therapy, brentuximab vedotin, to target tumor cells; however, anti-CD30 drugs could diminish PZA activity in TB co-infected patients and consequently promote PZA resistance in *Mtb* (48, 49).

Importantly, this study also provides an explanation for Walsh McDermott’s perplexing observation that PZA monotherapy often elicited bewildering and unpredictable results (1). Namely, particular attention should be given to the host’s immune status regarding successful treatment with PZA. PZA refractory results have been observed for some patients co-infected with HIV/AIDS (50-53). HIV/AIDS infection has been shown to diminish and/or abrogate host responses related to phagosome maturation, including acidification and the oxidative burst, in addition to low CD4 T cell numbers (54-57). However, these findings regarding refractory responses to PZA in HIV patients have been challenged by results reported by Chideya et al., which show lower PZA concentrations (*Cmax*) in patients co-infected with HIV (58). Thus, it is possible that PZA treatment fails in HIV co-infected patients due to suboptimal drug concentrations rather than a true refractory response. Studies should further seek to characterize PZA drug concentrations in HIV patients as well as immune deficient murine lines, such as the ones used in this study. Undoubtedly, understanding which patient populations that may be unresponsive to PZA treatment will enable physicians to better tailor TB drug regimens to ensure successful *Mtb* sterilization and prevent future relapses.

These findings also highlight several opportunities for host-directed therapies and other antitubercular drugs to either enhance PZA activity or overcome deficiencies in the host response. Adjunctive therapies may include those that induce the autophagy pathway and thereby promote phagosome maturation and the oxidative burst. For example, a different antitubercular drug, bedaquiline, increases PZA efficacy by promoting PZA accumulation in phagosomes by inducing lysosome acidification via autophagy (20, 59). Commercially available host-directed drugs, such as rapamycin and metformin, that also induce autophagy may potentiate PZA and should be further evaluated for repurposing in the anti-TB drug regimen (60-64). Additionally, the development of recombinant forms of CD153 will allow for CD30 stimulation and promote IFNγ independent CD4 T cells and other effector functions that drive PZA activity in the lungs. In a broader context, our data demonstrates the need to include the host during the initial stages of drug development. It is possible that several promising drugs are overlooked and eliminated when solely using *in vitro* methods that focus on the drug-parasite paradigm. Thus, it is essential that we restore the drug-parasite-host relationship for the consideration and evaluation of newly discovered antitubercular drugs and anti-TB combinatorial therapies.

## Materials and Methods

### Ethics Statement

Animal protocols used in this study were reviewed and approved by the University of Minnesota Institutional Animal Care and Use Committee (IACUC) under protocol 2407-42244A and Association for Assessment and Accreditation of Laboratory Animal Care, respectively. The University of Minnesota’s NIH Animal Welfare Assurance Number is D16-00288. All animal experiments were strictly conducted in accordance with the recommendations in the Guide for the Care and Use of Laboratory Animals of the National Institutes of Health (65).

### Mice

WT C57BL/6J (#000664), WT C57BL/6NJ (#005304), B6N.129S2-*Ncf1*^*tm1Shl*^/J (PHOX47^phox-/-^, #027331), B6129S7-*IfngR1*^*tm1Agt*^/J (IFNGR1^-/-^, #003288), and B6.129S2-*H2*^*dlAb1-Ea*^/J (MHCII^-/-^, #003584) were purchased from the Jackson Laboratory. Mice were 4-7 weeks of age. All animals were housed in an ABSL-3 vivarium except for mice used for bone marrow extraction. Mice were provided with irradiated standard rodent chow and sterilized water *ad libitum*..

### Cell Line and Primary Cells

CMG14-12 cells (C3H male) were cultured in Dulbecco’s Modified Eagle Medium/Nutrient Mixture F-12 (DMEM/F12) supplemented with 10% heat-inactivated fetal bovine serum (HI-FBS), and 1% penicillin/streptomycin mixture at 37°C in a 5% CO_2_ incubator (66). CMG14-12 conditioned medium was filtered through a 0.22 µm sterile syringe unit with a polyethersulfone membrane, aliquoted, and frozen at -70°C until further use. Macrophages were harvested separately from the bone marrow of female C57BL/6J, C57BL/6NJ, B6N.129S2-*Ncf1*^*tm1Shl*^/J, and B6129S7-*IfngR1*^*tm1Agt*^/J mice and maintained in DMEM/F12 supplemented with HI-FBS, 1% penicillin/streptomycin mixture, and 5% CMG14-12 conditioned medium. CMG14-12 conditioned medium was utilized as a source for macrophage colony-stimulating factor (M-CSF). Macrophages were washed thrice in Hank’s Buffered Saline Solution (HBSS) and seeded at 2.0 × 10^5^ cells/mL in 12 well plates. Designated macrophages were activated using recombinant interferon-gamma (IFN-γ; Fujifilm Irvine Scientific, 5ng/mL) for 18-24 h. Antibiotics were omitted from macrophage culture medium 24 h prior to and throughout *Mtb* infection.

### Bacterial Culture and Macrophage Infection

All experiments involving the use of *Mtb* were conducted inside a BSL-3 facility. *Mtb* H37Rv was used throughout this study. *Mtb* was grown in Middlebrook 7H9 (MB7H9) medium containing 0.5% glycerol, 10% oleic acid-dextrose-catalase (OADC; BD Bioscience) supplement, and 0.02% tyloxapol at 37°C with mild shaking (100 rpm). For macrophage infection, mid-logarithmic *Mtb* was washed thrice in 1X phosphate buffered saline (PBS) containing 0.05% Tween 80 (TW80) and resuspend as a single cell suspension in macrophage culture medium without antibiotics. Single cell suspension of the *Mtb* inoculum was achieved by allowing the prepared culture to incubate for 5 min at room temperature (RT) for any clumps to sediment. The top two-thirds of the *Mtb* suspension was used for macrophage infection. For infection, *Mtb* was added to macrophages at a multiplicity of infection (MOI) of 1. After 2 h of infection, extracellular bacteria were removed by washing macrophages thrice in HBSS (1.0 mL/well). Macrophage medium with/out pyrazinamide (200 and 400 µg/mL) was changed daily for the entire experiment. IFN-γ (5 ng/µL) was added every 2 days to designated cells. Infected macrophages were maintained in a 5% CO_2_ incubator at 37°C for 5 days. Macrophages were washed thrice in HBSS and lysed using 0.1% Triton-X solution at days 0, 1, 2, 4, and 5. Macrophage homogenates were serially diluted, plated on Middlebrook 7H10 agar supplemented with 0.5% glycerol and 10% OADC, and incubated at 37°C for 3 weeks until bacterial enumeration. Cell infection experiments were conducted in triplicate with two technical replicates for each treatment and timepoint.

### Mice Infection and Drug Treatment

Male and female C57BL/6J (n = 50), IFNGR1^-/-^ (n = 57), and MHC II^-/-^ (n = 50) mice were infected with ∼100 CFU of *Mtb* using an inhalation exposure system (Glas-Col) as previously described (67). Nine to 10 mice (n= 5 male and n= 4-5 female) from each strain were sacrificed by CO_2_ inhalation 24 h post-infection to determine the number of CFU established in the lungs. *Mtb* infection was allowed to progress for 3 weeks (pretreatment) to initiate the cell-mediated immune response and 9-10 mice (n = 5 male and n = 4-5 female) were euthanized as before and lungs, livers, and spleens were plated for bacterial enumeration. After the pretreatment, drug treatment occurred. Pyrazinamide (PZA; 150 mg/kg), isoniazid (INH; 30 mg/kg), and sterile water (vehicle control) were administered to designated groups (n=10-13 male and female per treatment and mouse strain) by oral gavage for 5 days a week for a total of 2 weeks. At completion of drug treatment, mice were euthanized by CO_2_ inhalation and organs were removed for bacterial enumeration.

### Enumeration of bacterial load

At the time of sacrifice, lungs, livers, and spleens were aseptically removed from mice. All organs were mechanically disrupted in 1X PBS-Tw80 using a tissue homogenizer. Organ homogenates were serially diluted and plated on MB7H10 agar supplemented with 0.5% glycerol, 10% OADC, and cycloheximide (10 mg/mL). Colonies were enumerated after at least 21 days of incubation at 37°C. Total bacterial burden was calculated based on organ volume.

### Statistical Analysis

Comparisons of *in vivo* treatment groups at specified timepoints were analyzed using paired two-tailed *t-*tests and *p* values were adjusted for multiple comparisons. Two-way ANOVA with Bonferoni correction was employed for BMDM infection assays. Graphpad Prism software ver. 10.2.2 was used to construct figures.

## Acknowledgements

The authors would like to thank Dr. Tyler D. Bold, University of Minnesota, for his generous gift of CMG14-12 cells. The authors thank the University of Minnesota’s Biosafety Level 3 Program and the Research and Animal Resources area staff. This research was supported by the NIH R01 AI123146 awarded to ADB. EAL was supported by the NIH diversity supplement award R01AI123146-03S1.

## Supplementary Data

**Table S1:** Log differences in statistically significant treatment comparisons in *Mtb* infected C57BL6/J mice BMDMs.

| **Day 4** |  |  |  |
| --- | --- | --- | --- |
| **C57BL6/J** | **Comparison** | **Log<sub>10</sub> Difference** | **P -value** |
|  | PZA<sub>400</sub> v. ND | 0.912 | 0.0278 |
| | IFN-γ v. ND | 1.51 | 0.0201 |
| | IFN-γ + PZA<sub>200</sub> v. ND | 2.11 | 0.0184 |
| | IFN-γ + PZA<sub>400</sub> v. ND | 2.77 | 0.0180 |
|  | PZA<sub>400</sub> v. PZA<sub>200</sub> | 0.782 | 0.035 |
| | IFN-γ v. PZA<sub>200</sub> | 1.38 | 0.0223 |
| | IFN-γ v. PZA<sub>400</sub> | 0.6 | 0.05 |
| | IFN-γ + PZA<sub>200</sub> v. PZA<sub>200</sub> | 1.977 | 0.0199 |
| | IFN-γ + PZA<sub>400</sub> v. PZA<sub>200</sub> | 2.64 | 0.0013 |
| | IFN-γ + PZA<sub>200</sub> v. PZA<sub>400</sub> | 1.19 | 0.0187 |
| | IFN-γ + PZA<sub>400</sub> v. PZA<sub>400</sub> | 1.86 | 0.0192 |
| | IFN-γ + PZA<sub>400</sub> v. IFN-γ | 1.26 | 0.0189 |
| | IFN-γ + PZA<sub>400</sub> v. IFN-γ + PZA<sub>200</sub> | 0.663 | 0.05 |
| **Day 5** |  |  |  |
|  | **Comparison** | **Log<sub>10</sub> Difference** | **P -value** |
|  | PZA<sub>400</sub> v. ND | 1.23 | 0.0079 |
| | IFN-γ v. ND | 2.22 | 0.0041 |
| | IFN-γ + PZA<sub>200</sub> v. ND | 3.02 | 0.0038 |
| | IFN-γ + PZA<sub>400</sub> v. ND | 3.60 | 0.0031 |
|  | PZA<sub>400</sub> v. PZA<sub>200</sub> | 0.868 | 0.0345 |
| | IFN-γ v. PZA<sub>200</sub> | 1.85 | 0.0190 |
| | IFN-γ v. PZA<sub>400</sub> | 0.984 | 0.0269 |
| | IFN-γ + PZA<sub>200</sub> v. PZA<sub>200</sub> | 2.65 | 0.0178 |
| | IFN-γ + PZA<sub>400</sub> v. PZA<sub>200</sub> | 3.24 | 0.0034 |
| | IFN-γ + PZA<sub>200</sub> v. PZA<sub>400</sub> | 1.78 | 0.0188 |
| | IFN-γ + PZA<sub>400</sub> v. PZA<sub>400</sub> | 2.37 | 0.0045 |
| | IFN-γ + PZA<sub>200</sub> v. IFN-γ | 0.8 | 0.0322 |
| | IFN-γ + PZA<sub>400</sub> v. IFN-γ | 1.38 | 0.0082 |
| | IFN-γ + PZA<sub>400</sub> v. IFN-γ + PZA<sub>200</sub> | 0.585 | 0.0356 |

**Table S2:** Log differences in statistically significant treatment comparisons in *Mtb* infected C57BL6/NJ mice BMDMs.

| **Day 4** |  |  |  |
| --- | --- | --- | --- |
| **C57BL6/NJ** | **Comparison** | **Log<sub>10</sub> Difference** | **P -value** |
|  | PZA<sub>400</sub> v. ND | 0.692 | 0.0394 |
| | IFN-γ v. ND | 1.04 | 0.0244 |
| | IFN-γ + PZA<sub>200</sub> v. ND | 1.33 | 0.0201 |
| | IFN-γ + PZA<sub>400</sub> v. ND | 1.85 | 0.0174 |
|  | PZA<sub>400</sub> v. PZA<sub>200</sub> | 0.360 | 0.0031 |
| | IFN-γ v. PZA<sub>200</sub> | 0.709 | 0.0003 |
| | IFN-γ + PZA<sub>200</sub> v. PZA<sub>200</sub> | 1 | 0.0003 |
| | IFN-γ + PZA<sub>400</sub> v. PZA<sub>200</sub> | 1.54 | 0.0002 |
| | IFN-γ + PZA<sub>200</sub> v. PZA<sub>400</sub> | 0.64 | 0.0461 |
| | IFN-γ + PZA<sub>400</sub> v. PZA<sub>400</sub> | 1.18 | 0.0211 |
| | IFN-γ + PZA<sub>400</sub> v. IFN-γ | 0.831 | 0.0032 |
| | IFN-γ + PZA<sub>400</sub> v. IFN-γ + PZA<sub>200</sub> | 0.54 | 0.0010 |
| **Day 5** |  |  |  |
|  | **Comparison** | **Log<sub>10</sub> Difference** | **P -value** |
|  | PZA<sub>400</sub> v. ND | 0.953 | 0.0427 |
| | IFN-γ v. ND | 1.34 | 0.0322 |
| | IFN-γ + PZA<sub>200</sub> v. ND | 1.87 | 0.0282 |
| | IFN-γ + PZA<sub>400</sub> v. ND | 2.77 | 0.0269 |
|  | PZA<sub>400</sub> v. PZA<sub>200</sub> | 0.6 | 0.0077 |
| | IFN-γ v. PZA<sub>200</sub> | 0.985 | 0.0034 |
| | IFN-γ v. PZA<sub>400</sub> | 0.389 | 0.0003 |
| | IFN-γ + PZA<sub>200</sub> v. PZA<sub>200</sub> | 1.51 | 0.0025 |
| | IFN-γ + PZA<sub>400</sub> v. PZA<sub>200</sub> | 2.41 | 0.0022 |
| | IFN-γ + PZA<sub>200</sub> v. PZA<sub>400</sub> | 0.914 | <0.0001 |
| | IFN-γ + PZA<sub>400</sub> v. PZA<sub>400</sub> | 1.82 | <0.0001 |
| | IFN-γ + PZA<sub>200</sub> v. IFN-γ | 0.524 | 0.0090 |
| | IFN-γ + PZA<sub>400</sub> v. IFN-γ | 1.43 | 0.0033 |

